# A calibration and uncertainty quantification analysis of classical, fractional and multiscale logistic models of tumour growth

**DOI:** 10.1101/2023.04.12.536622

**Authors:** Nikolaos M. Dimitriou, Ece Demirag, Katerina Strati, Georgios D. Mitsis

## Abstract

The validation of mathematical models of tumour growth is frequently hampered by the lack of sufficient experimental data, resulting in qualitative rather than quantitative studies. Recent approaches to this problem have attempted to extract information about tumour growth by integrating multiscale experimental measurements, such as longitudinal cell counts and gene expression data. In the present study, we investigated the performance of several mathematical models of tumour growth, including classical logistic, fractional and novel multiscale models, in terms of quantifying *in-vitro* tumour growth in the presence and absence of therapy. We further examined the effect of genes associated with changes in chemosensitivity in cell death rates. State-of-the-art Bayesian inference, likelihood maximisation and uncertainty quantification techniques allowed a thorough evaluation of model performance. The results suggest that the classical single-cell population model (SCPM) was the best fit for the untreated and low-dose treatment conditions, while the multiscale model with a cell death rate symmetric with the expression profile of OCT4 (Sym-SCPM) yielded the best fit for the high-dose treatment data. Further identifiability analysis showed that the multiscale model was both structurally and practically identifiable under the condition of known OCT4 expression profiles. Overall, the present study demonstrates that model performance can be improved by incorporating multiscale measurements of tumour growth.

## 1. Introduction

Over the past few decades, mathematical models have provided significant insights into the understanding of tumour growth and its underlying mechanisms [1, 2, 3, 4, 5, 6, 7, 8, 9, 10, 11, 12, 13, 14, 15, 16], as well as the optimization of treatment [17, 18, 19]. One of the goals of the emerging field of mathematical oncology is the translation of models from an experimental to a clinical setting, contributing to personalized therapy [17, 20, 21, 22, 23]. In this context, computational and experimental challenges have shifted from the qualitative and informative use of models to patient-specific predictions.

A major computational challenge is the selection of an appropriate mathematical model. Tumour growth is a complex phenomenon that can be modelled at multiple levels. At the macroscopic level, several models can provide a longitudinal description of tumour growth, including exponential-linear, Gompertz, Verhulst or logistic, and von Bertalanffy models [24] and, more recently, the expansion of these models to incorporate additional cell populations, such as drug sensitive and resistant cells [18]. Longitudinal measurements of tumour growth have been used to calibrate mathematical models. Depending on the cancer type, case and stage, some models may be more appropriate than others [2]. Recent studies have focused on several aspects that make a model suitable for a specific cancer type and case. These include the degree to which a model can describe the tumour behaviour, and the uncertainties associated with the model prediction [25]. Similarly, studies have focused on evaluating of methods for estimating model parameters from experimental data [26, 27]. Although these techniques can provide a robust assessment of model predictions, this is only possible when appropriate experimental data are available.

The sparsity of experimental measurements and single-level data extraction may limit the ability of a model to accurately describe tumour growth. Recent studies have addressed this problem by incorporating multiscale data to describe tumour growth. Johnson et al. [28] integrated longitudinal bulk cell count measurements and single-cell transcriptomic data to estimate the ratio between drug-sensitive and drugresistant cell populations at multiple time points, resulting in more accurate model predictions. Similarly, Smalley et al. [19] used single-cell mRNA analysis to define the transcriptional heterogeneity of melanoma and its response to BRAF inhibitors. They combined these data with a mathematical model for maintaining competition between drug-sensitive and drug-resistant states to control resistance.

Promising evidence for the use of gene expression data integrated with bulk cell count measurements has emerged from studies of bacterial growth [29]. Klumpp et al. [30] demonstrated specific patterns between gene expression levels and growth rates. Studies in *E. coli* [31, 32] have shown that the RNA/protein ratio is linearly correlated with the specific growth rate *λ* = (ln 2)*/*doubling time. Although bacteria are much simpler organisms than cancer cells, it is not unreasonable to assume that a similar phenomenon may occur in cancer [33], specifically between cell death rates and the expression of genes involved in altering chemosensitivity, such as OCT4 [34, 35, 36].

In the present study, we provide a thorough investigation of classical, fractional and multiscale models of tumour growth. We calibrated these models against experimental measurements of cell counts from the C33A cell line. We used two calibration methods: likelihood optimization using the Improved Stochastic Ranking Evolution Strategy (ISRES) algorithm [37] and Bayesian inference using a Transitional Markov Chain Monte Carlo algorithm [38]. The multiscale models incorporated information on both cell counts and gene expression data possibly contributing to the change in chemosensitivity. We examined two experimental conditions; no treatment, and treatment with 5FU. We found that the single-cell population model (SCPM) was the most appropriate model for non-treatment and low-treatment conditions, exhibiting strong agreement with the experiments and low uncertainty compared to the other models. For higher doses, we investigated four multiscale models derived from the classical SCPM and the fractional model (FM). We integrated these models with death rate parameters that took into account gene expression profiles based on a gene expression model. The multiscale model with a death rate symmetrical to the expression profile of OCT4 (Sym-SCPM) showed a significant improvement in terms of agreement with the data and uncertainty. Since this model contains a novel cell death term, we found necessary to examine whether the model parameters can be determined by the model outputs (i.e. structural identifiability) and estimate their uncertainties given the presented data (i.e. practical identifiability) [39]. Our analysis concluded that the Sym-SCPM is both structurally and practically identifiable, and showed the important role of gene expression data with regards to the latter.

## 2. Methods

### 2.1. Experiments

Cell culture, 5FU treatment and counting: C33A cells were obtained from ATCC and cultured in MEM medium, 1% Penicillin/Streptomycin, 1% L-Glutamine and 10% FBS. The cell media were regularly monitored using PCR with mycoplasma-specific primers to ensure that the cultures remained mycoplasma-negative throughout the experiment. Fresh 5FU (Sigma Aldrich) was prepared weekly. 5 × 10^4^ C33a cells were plated in 24-well plates for cell counting, and 1 × 10^6^ cells were plated in 6-well plates for RNA extraction. One plate was prepared for each time point. Each biological replicate consisted of three technical replicates per treatment condition. Three days post-seeding, the cells were treated with 100, 10, 1, 0.1 and 0.01 μM 5FU diluted in media and incubated indicated time-points. Cell counts were performed using a hemocytometer, and dead cells were omitted from counting using trypan blue exclusion.

RNA Extraction and qRT-PCR RNA extraction was performed using Qiagen RNeasy Mini Kit (169014761) according to the manufacturer’s instructions. RNA yield and quality (260 nm/280 nm ratio) were measured using Nanodrop™. RNA integrity was further validated on 1% agarose gel (28S and 18S rRNA bands). cDNA was synthesized using iScript cDNA synthesis kit, according to the manufacturer’s instructions. Following cDNA synthesis, qRT-PCR was performed using the KAPA SYBR® FAST qPCR Master Mix (2x) kit. PCR was performed on a CFX96 qPCR system (BioRad). The following primer sequences were used (Oct4: Fwd: GAAGGATGTGGTCCGAGTGT, Rev: GTGAAGTGAGGGCTCCCATA, GAPDH: Fwd: TGCACCACCAACTGCTTAGC, Rev: GGCATGGACTGTGGTCATGAG). Relative expression levels were determined using the DCT method.

### 2.2. Classical mathematical models

The *classical single-cell population model* (1) is a logistic growth model with three parameters; growth rate *r*_*g*_, carrying capacity *K*, and effective cell death rate *u*_*eff*_. The variable *N* = *N* (*t*) represents the number of cancer cells over time. The effective cell death rate depends on the drug dose *u* and the death rate *λ* hence, *u*_*eff*_ = *λu*.

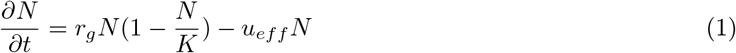

The *two-cell population model* considers both drug-sensitive and drug-resistant cancer cells. The model in (2)-(4) is similar to the single-cell population model found in (1) with the addition of the parameter *α*, which represents the spontaneous transition rate of sensitive cells to resistant cells. An additional constraint is that the effective death rate of sensitive cells, *u*_*eff,s*_, should be greater than the effective death rate of resistant cells, *u*_*eff,r*_, i.e., *u*_*eff,s*_ *< u*_*eff,r*_. Similarly, the effective death rates depend on the drug dose and the death rate parameter, and they are equal to *u*_*eff,s*_ = *λ*_*s*_*u, u*_*eff,r*_ = *λ*_*r*_*u*. Since the initial numbers of sensitive and resistant cells were unknown, we considered unknown initial conditions for the two populations. The total initial number of cells was determined from the experiments; hence, we chose to estimate the resistant and sensitive cells at *t* = 0 from (4), where *T* is the total number of cells.

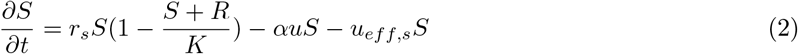

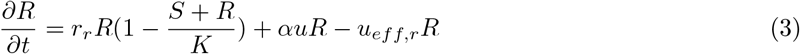

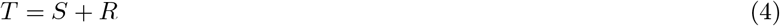

Both models were numerically solved using the 5-order accurate Runge-Kutta Dormand-Prince method coded in C++ [40] and the 5-order accurate Tsitouras method in Julia [41, 42].

### 2.3. Fractional mathematical model

We examined the generalized fractional version of the single-cell population model as found in (1). The fractional logistic model defined in (5) takes into account the growth rate *r*_*g*_, carrying capacity *K*, and the order of the derivative *α*. The order of the derivative takes the values 0 *< α* ≤ 1 and is considered as unknown parameter during model calibration. The model was solved analytically using the formulation of Caputo’s derivatives, the Mittag-Leffler function, and the Laplace transform (Appendix).

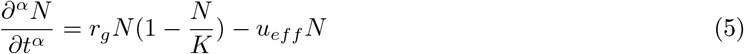

### 2.4. Multiscale classical and fractional models

Previous studies have shown that gene expression and cell proliferation can be interdependent [29]. In this section, we provide a multiscale formulation of classical and fractional models by combining gene expression levels and cell counts in a single equation.

#### 2.4.1. Correlation of cell counts with gene expression profiles

The OCT4, KLF4, SOX2, and NANOG genes likely play an important role in the response of cancer cells to chemotherapy [34, 35, 36]. We used longitudinal data of the expression profiles of these genes obtained by real-time quantitative Polymerase Chain Reaction (RT-qPCR) to quantify the relation between expression profiles and cell counts. To do this, we normalized the cell counts of the treatment data with respect to the control data and then estimated the Pearson correlation coefficient [43] between the normalized cell counts and the gene expression profiles.

#### 2.4.2. Model integration with the effects of gene expression in cell death

For the multiscale expansion of the models presented above, we used the expression of genes that exhibited the strongest association with cell counts. These profiles were then fitted to a simple asymptotic model described by

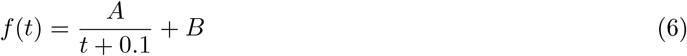

using the least-squares method. The gene expression model was incorporated into the cell death parameter of the corresponding classical single-population model, as well as the order of the derivative for the fractional model.

Since the increase in the expression of genes such as OCT4 has been associated with chemoresistance and, in turn, a lower cell death rate [34, 35, 36], we examined two possible ways of relating the cell death term with the gene expression model. One of the examined cell death parameter is the *inverted* gene expression model of the form

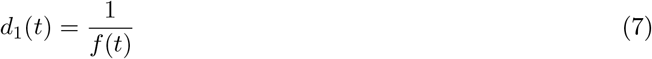

Second, we considered the cell death parameter to be *symmetric* with respect to the OCT4 expression, i.e.,

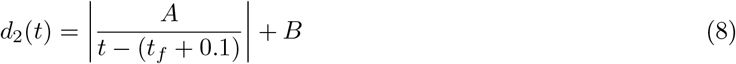

where *t*_*f*_ denotes the final time point of the experiment. We introduced these time-dependent cell death rates into the cell death term of the classical single cell population model,

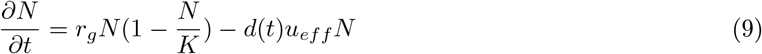

where *d*(*t*) = *d*_1_(*t*) or *d*(*t*) = *d*_2_(*t*). Additionally, we considered coupling the order of the fractional model with the cell death rates as follows,

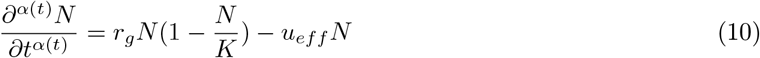

where *α*(*t*) = *d*(*t*)*/* max{*d*(*t*)} because of the constraint 0 *< α* ≤ 1 imposed on the fractional model solution.

### 2.5. Calibration

In this section, we describe the two methods used to calibrate the model parameters with respect to the cell count data. The first is a maximum likelihood optimization method and the second is Bayesian inference for estimating the Posterior Distribution Functions (PDFs) of the model parameters. We continue with the calibration strategies that allowed us to assess the contribution of the model parameters to the model outputs.

#### Maximum Likelihood estimation

We used the Improved Stochastic Ranking Evolution Strategy (ISRES) algorithm [37] found in the NLopt package [44] in Julia [45]. Similar to any Evolutionary Strategy algorithm, ISRES uses an array of candidate solutions that are updated based on *mutation* and *selection* rules. Mutation represents the step size between the parent solution and the offspring. In ISRES, mutation is controlled via a log-normal step size update and exponential smoothing. To avoid biases introduced by spherical symmetry assumptions [46], the ISRES algorithm performs a differential variation step using a Nelder-Mead-like update method [47, 48]. The selection rule is based on fitness ranking, and in this study, the fitness ranking was the log-likelihood function. The likelihood was defined as

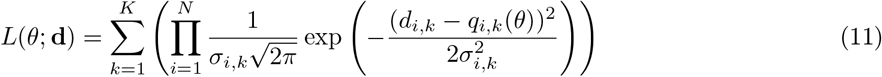

where *N* is the total number of time points, *K* is the number of datasets, **d** the cell count, *d*_*i,k*_, *q*_*i,k*_ the cell count of the experimental sample and simulation result, respectively, at time point *i* and dataset *k*, and *σ* _*i,k*_ is the variance of the distribution of the likelihood obtained from the experimental error. Here, *σ* _*i,k*_ was the standard deviation of the three measurements for each time point for all datasets. The number of candidate solutions at each step was 20 × (*n* + 1), where *n* is the number of model parameters.

#### Bayesian inference

We used the PDFs of the model parameters and the counts from the cell culture data, *D*, to estimate the most probable parameter values using Bayes’ rule

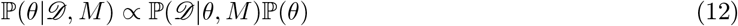

where ℙ(*θ*|*D, M*) is the posterior PDF of the model parameters *θ* given the observed data *D* and the model *M*, ℙ(*D*|*θ, M*) is the likelihood of the observed data *D* given the model *M* and parameters *θ*, and ℙ(*θ*) is the prior PDF. We assumed uninformative, uniform distributions for the model parameter prior PDFs. The experimental data consisted of three replicates, each containing samples from 10 time points. The datasets were then combined to estimate the likelihood (Eq. (11)).

We used a Transitional Markov Chain Monte Carlo (TMCMC) algorithm implemented in the Π4U package [38]. The TMCMC algorithm iteratively constructs series of intermediate posterior PDFs as follows,

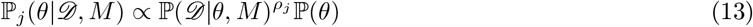

where *j* = 0, …, *m* is the Monte Carlo time series index (generation index), *ρ*_*j*_ controls the transition between generations, and 0 *< ρ*_0_ *< ρ*_1_ *<* … *< ρ*_*m*_ = 1. The TMCMC method uses a large number of parallel chains that are evaluated at each step to produce a result that is close to the true posterior PDF.

#### Calibration strategies

The increasing complexity of the selected models allowed us to follow an incremental calibration strategy. Specifically, the calibration of the single-population model of (1) without treatment, i.e. *u*_*eff*_ = 0, allowed us to estimate the growth rate, *r*_*g*_, and the carrying capacity, *K*. For the treatment datasets, instead of considering these two parameters as completely unknown, we used a uniform prior bounded by the 95% credible intervals (CI), calculated using the high-density interval method [49], of the posterior distributions obtained from the calibration on the control datasets. This process was repeated for all the models examined, as shown in Table 1.

**Table 1:** Prior distributions of the model parameters. The employed calibration strategy assumes that the growth rates and carrying capacities are not affected by the treatment. Thus, they can be estimated from the control (no treatment) data. The *U* notation denotes uniform distribution, and the *U* [*CI*_95_] denotes uniform distribution with boundaries the 95% credible interval of parameter posterior distribution found from the control studies. The subscripts *s* and *r* correspond to sensitive and resistant populations, respectively.

| Model | Priors |  |  |  |  |  |  |
| --- | --- | --- | --- | --- | --- | --- | --- |
| | $r_g$ | $r_{s,r}$ | $K$ | $\alpha$ | $u_{eff}$ | $u_{eff,s,r}$ | $R_0$ |
| SCPM-C | $U[0, 0.15]$ | - | $U[3, 5] \cdot 10^5$ | - | - | - | - |
| SCPM-T | $U[CI_{95}]$ | - | $U[CI_{95}]$ | - | $U[0, 0.15]$ | - | - |
| TCPM-C | - | $U_s[0.05, 0.15]$<br>$U_r[0.01, 0.05]$ | $U[3, 5] \cdot 10^5$ | - | - | - | $U[0, 5 \cdot 10^5]$ |
| TCPM-T | - | $U_s[CI_{95}]$<br>$U_r[CI_{95}]$ | $U[CI_{95}]$ | $U[0, 0.2]$ | - | $U_s[0.0011, 0.1]$<br>$U_r[0.0, 0.0010]$ | $U[CI_{95}]$ |
| FM-C | $U[0, 0.15]$ | - | $U[3, 5] \cdot 10^5$ | $U(0, 1)$ | - | - | - |
| FM-T | $U[CI_{95}]$ | - | $U[CI_{95}]$ | $U(0, 1)$ | $U[0, 0.15]$ | - | - |
| Inv/Sym-SCPM-C | $U[0, 0.15]$ | - | $U[3, 5] \cdot 10^5$ | - | - | - | - |
| Inv/Sym-SCPM-T | $U[CI_{95}]$ | - | $U[CI_{95}]$ | - | $U[0, 0.15]$ | - | - |
| Inv/Sym-FM-C | $U[0, 0.15]$ | - | $U[3, 5] \cdot 10^5$ | $U(0, 1)$ | - | - | - |
| Inv/Sym-FM-T | $U[CI_{95}]$ | - | $U[CI_{95}]$ | $U(0, 1)$ | $U[0, 0.15]$ | - | - |

To assess the quality of the fits and the agreement between the simulation and the experimental data, we calculated the Concordance Correlation Coefficient (CCC) [50], defined as

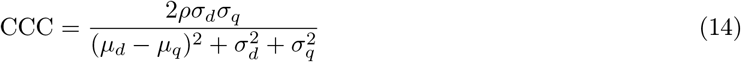

where *n* is the number of time points in each dataset, *ρ* is the Pearson correlation coefficient [43], and 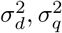 are the variances of the *in-vitro* and *in silico* data, respectively.

### 2.6. Uncertainty analysis

The use of Bayesian inference and the TMCMC algorithm allows estimation of the posterior density of the model parameters. To assess the effect of parameter uncertainty on the resulting cell counts, we performed 5000 simulations of each model with random sets of parameters drawn from their corresponding posterior distributions. The parameters were selected from a unknown distribution using the inverse transform sampling technique [51].

### 2.7. Identifiability of model parameters

Model identification falls into two categories: structural and practical. Structural identifiability examines the identification of model parameters by assuming an infinite number of error-free data points. On the other hand, practical identifiability examines the possibility of uniquely identifying the model parameters given finite and noisy sets of observations. This section examines the structural and practical identifiability of the best candidate model. For structural identifiability, we followed a methodology similar to that of [52, 18, 28], which includes the use of Lie derivatives.

Simpson et al. [53] compared two methods to test the practical identifiability of a biophysical model using Bayesian inference [54] and the profile likelihood method [55]. The two methods gave similar results. As Bayesian inference yields full posterior distributions, we used it to examine the practical identifiability of the best candidate model. Although we performed Bayesian inference during the calibration process, the prior distribution boundaries were set to be the *U* [CI_95_] obtained from the control data. In other words, we set as prior range to be the 95% credible interval of the posterior distributions of the parameters obtained from the calibration with the control data. Instead, for the identifiability analysis, we expanded the prior boundaries beyond these limits, and we set them equal to the prior boundaries originally used for the calibration with the control data.

## 3. Results

### 3.1. Classical single cell population model calibration

The calibration results of the single-cell population model for the control and treatment datasets are shown in Fig. 2. The posterior PDF manifold of the model parameters for the control data in Fig. 2g shows low uncertainty in the growth rate, *r*_*g*_, and a skewed posterior PDF towards the upper limit of the carrying capacity, *K*. The simulation results for the control (no treatment) data are shown in Fig. 2a and were in good agreement with the experimental data, while the uncertainty in the simulation output remained low. The good agreement between the experimental and simulation data can be confirmed by the CCC results in Table 3. Similar behaviour was observed for calibration with the treatment data (Fig. 2b-2f), where the parameters *r*_*g*_ and *K* were sampled around the posterior 95% CI obtained from the calibration of the control datasets. The posterior manifold for the highest dose calibration is shown in Fig. 2h and shows uniform distributions within the bounds obtained using the 95% CI of the control posterior PDF. The treatment parameter *u*_*eff*_ exhibits low uncertainty compared to the prior boundaries (Table 1).

**Figure 1:**
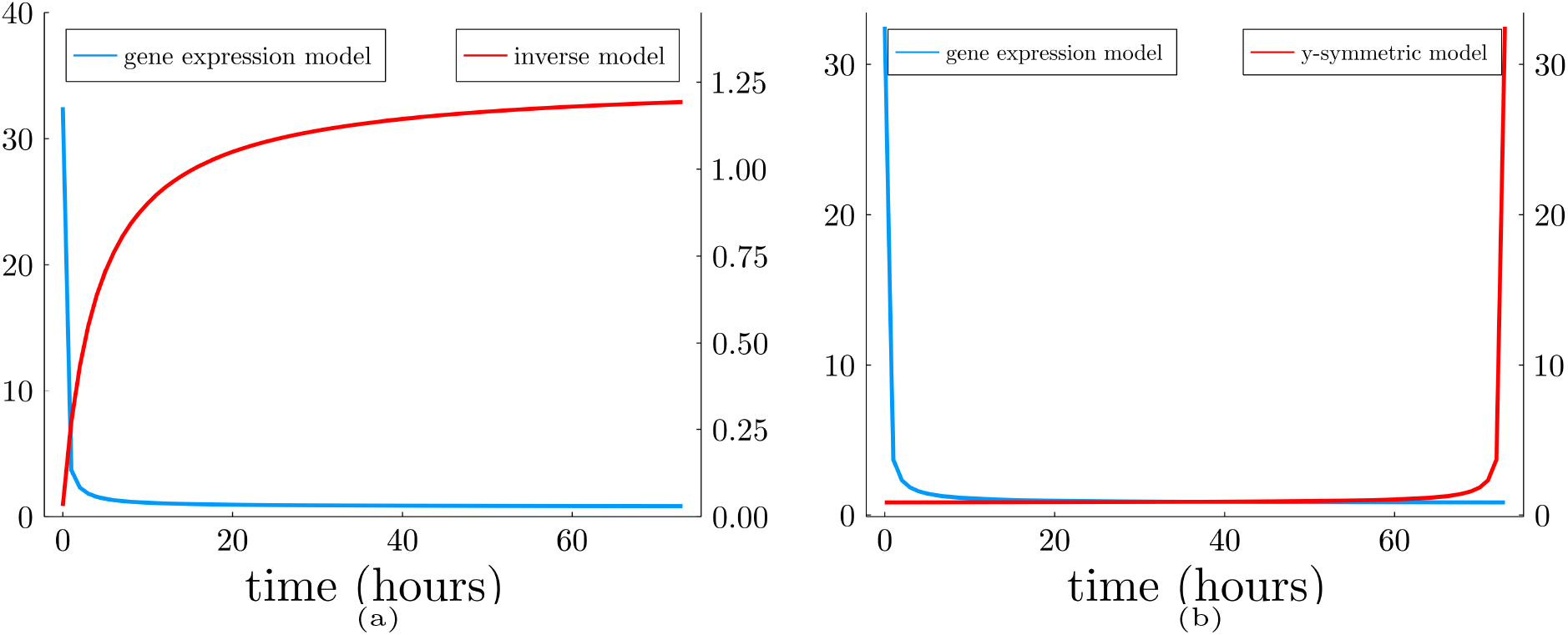
OCT4 expression model and cell death rates. (a) Inverted cell death rate (Eq. (7)), and (b) *y*-symmetric and shifted cell death rate (Eq. (8)) with respect to the gene expression.

**Figure 2:**
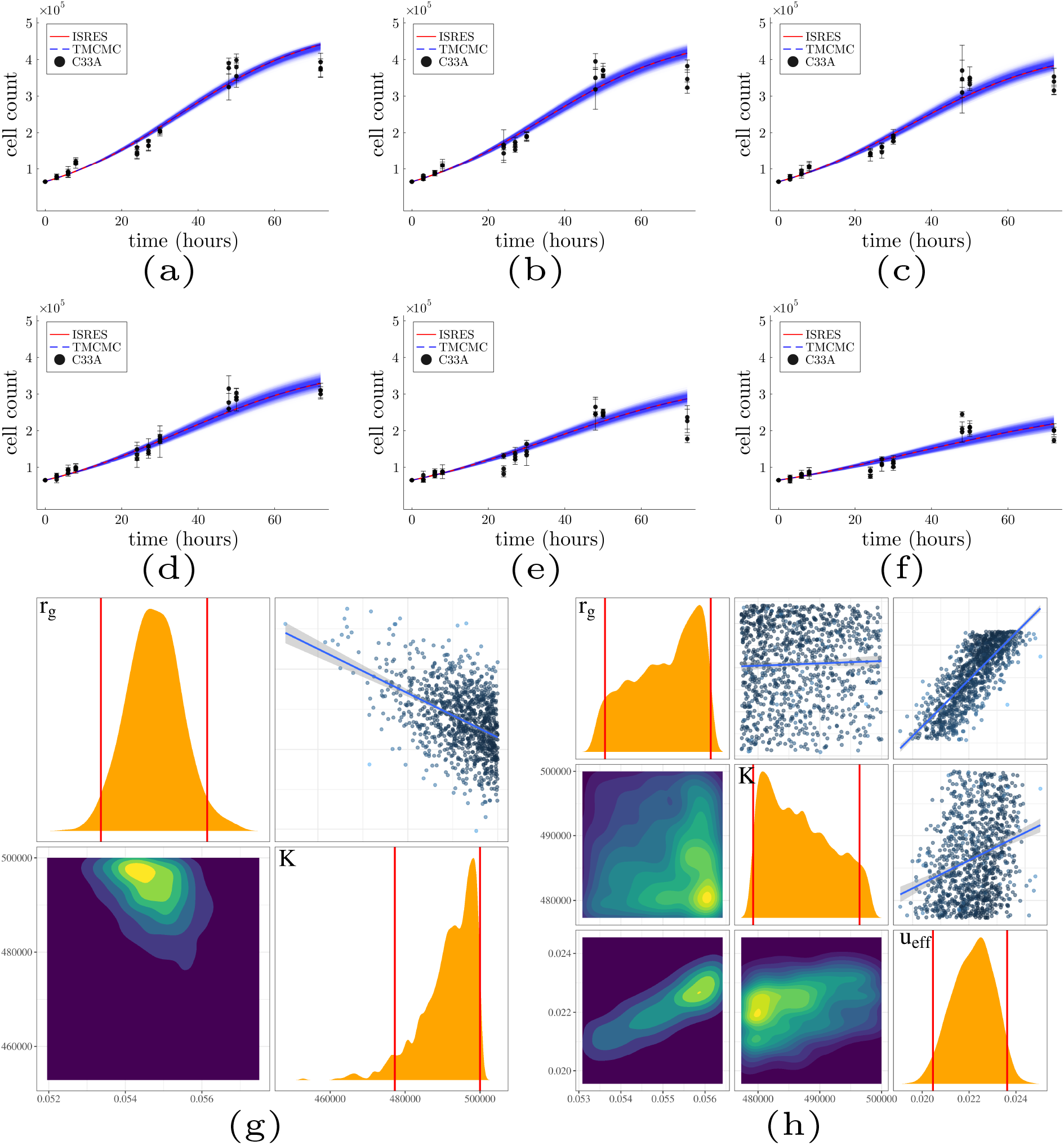
Calibration and uncertainty quantification of the classical single cell population model. (a) Control, (b) 0.01 μM, (c) 0.1 μM, (d) 1 μM, (e) 10 μM, and (f) 100 μM of 5FU. Posterior distributions of the model parameters for (g) control, (h) 100 μM of 5FU datasets. The prior boundaries of the *r*_*g*_ and *K* parameters of the treatment model are extracted from the 95% CI of the posterior distributions of the corresponding parameters of the control model. The red lines denote the 95% CI of the posterior distributions.

### 3.2. Classical two-cell population model calibration

Although the presence of resistance is not clear in these datasets, we examined the fitness of the two-cell population model, which takes into account drug-sensitive and drug-resistant cells. The calibration results for the two-cell population model are shown in Fig. A.8. The overall fitness result of the model is comparable to that of the single-cell population model. The posterior PDF manifold of the model parameters for the control data in Fig. A.8g shows increased uncertainty compared to the single-population model, particularly in the resistant growth rate parameter and the initial number of resistant cells. This increased uncertainty is reflected in the total number of cancer cells, as shown in Fig. A.8a-A.8f. The uncertainty in the initial number of resistant cells remained high at the highest dose, as shown in Fig. 2h.

### 3.3. Fractional model calibration

We generalized the classical single-cell population model by considering the derivative order as unknown *α* ∈ (0, 1]. The fractional model calibration yielded similar performance to that of the two-cell population model, as shown in Fig. A.9. The introduction of the derivative order as a variable resulted in increased uncertainty compared to the classical single-population model. In addition, the derivative order tended to converge to 1, suggesting that the single-cell population ODE model is more appropriate than the fractional model.

### 3.4. Multiscale model calibration

We performed a multiscale integration of macroscopic single-cell population models with the effects of gene expression. This step involved searching for patterns between cell death and gene expression changes in OCT4, SOX2, KLF4, and NANOG. To do this, we normalized the cell counts of treatment data with respect to the counts in the non-treatment data and calculated the Pearson correlation coefficient between the normalized cell counts and the gene expression profiles. The correlation of the expression profile of OCT4 with the corresponding cell counts increased with respect to the 5FU dose and showed a strong correlation at 10 and 100 μM doses of 5FU, as shown in Table 2 and Fig. 3a, 3b. We used the expression profiles obtained at 10 and 100 μM to generate two candidate models of gene expression. The expression of OCT4 decreased with time, as shown in Fig. 3c, 3d. Fitting the gene expression model of (6) gave *A* = 3.17 ± 0.49, *B* = 0.795 ± 0.076 for 100 μM 5FU, and *A* = 1.04 ± 0.27, *B* = 0.811 ± 0.042 for 10 μM 5FU.

**Table 2:** Pearson correlation coefficient between the normalized cell counts of the treatment data with respect to the controls, and the expression profiles of the genes OCT4, KLF4, SOX2, NANOG. The correlation of the normalized counts with respect to the expression of OCT4 increased as a function of dose, resulting in a strong correlation for the two highest doses.

| 5FU ($\mu\text{M}$) | OCT4 | KLF4 | SOX2 | NANOG |
| --- | --- | --- | --- | --- |
| 0.01 | -0.11 | 0.20 | 0.12 | -0.26 |
| 0.1 | 0.39 | 0.19 | 0.79 | 0.00 |
| 1 | 0.34 | -0.57 | 0.04 | -0.25 |
| 10 | 0.89 | -0.07 | 0.68 | 0.03 |
| 100 | 0.90 | -0.24 | 0.97 | 0.68 |

**Figure 3:**
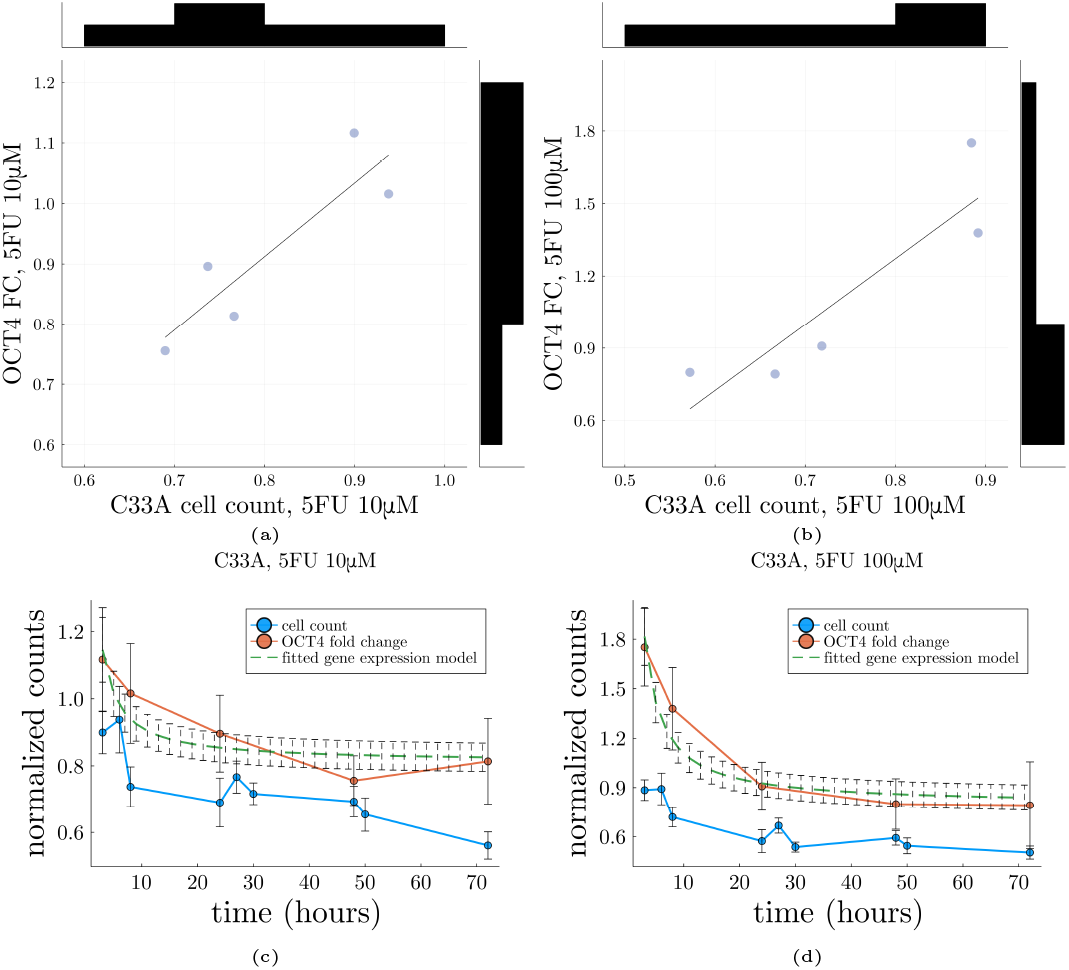
OCT4 expression and cell counts. Correlation of OCT4 expression profiles with normalized cell counts for (a) 10μM and (b) 100μM of 5FU. Gene expression model fitting results for (c) 10μM and (d) 100μM of 5FU. The error bars denote the standard deviation of the two model outputs.

Since the decrease in the expression of OCT4 has been associated with increased chemosensitivity [34, 35, 36], we examined two functions related to cell death. The first function takes the inverse of the gene expression model, and relates it to the cell death rate (7). The second function takes the *y*-axis symmetric and time-shifted gene expression model (8), and relates it to the cell death rate. We then combined the resulting time-varying cell death rates with two macroscopic single-cell population models (classical and fractional). For the classical model, time-varying cell death rates were introduced into the cell death term, resulting in two multiscale classical models (9) (namely Inv-SCPM and Sym-SCPM for coupling with the inverse and the *y*-symmetric gene expression models, respectively). For the fractional model, we introduced the time-varying cell death rates as a derivative order after normalizing them, resulting in two multiscale fractional models (10) (namely Inv-FM and Sym-FM for coupling with the inverse and the *y*-symmetric gene expression models, respectively).

The calibration results for the multiscale models are shown in Fig. 4, A.10. Overall, the multiscale SCPM performed better than the FM. The Sym-SCPM yielded the best performance among all models, achieving a curvature that the single-scale models were unable to capture at the two highest doses during the last days of the experiment Fig. 5. The sym-SCPM trajectories are more plausible than the corresponding classical SCPM because we expect cell elimination after prolonged times of the cell cultures, while the classical SCPM predicts stabilization of the population. The agreement with the experimental data, shown in Table 3, confirms that the Sym-SCPM exhibited the best performance. In contrast, the Sym-FM showed poor agreement with the experiments. Inv-SCPM and Inv-FM yielded similar performance to macroscopic models. The prior distribution boundaries of the parameters *r*_*g*_ and *K* were taken from the 95% CI posterior PDFs of the corresponding macroscopic no-treatment models, since the multiscale expansion focuses on cell death. The uncertainty of the multiscale model output remained low and was comparable to that of the macroscopic models. The derivative order scaling parameter *α* exhibited increased uncertainty for both Inv-FM and Sym-FM, as shown in Fig. A.10e and A.10f, suggesting insensitivity to the derivative scaling.

**Table 3:** Concordance correlation coefficient values between experimental data and mathematical models The abbreviated names SCPM, TCPM, FM denote single cell population model, two cell population model, and fractional model, respectively. The Inv and Sym abbreviations denote the integration of the corresponding model with the inverse gene model and *y*-symmetric model, respectively. The classical models performed better in no-treatment and low dose conditions, while the multiscale Sym-SCPM exhibited improved performance in high doses.

| Model | CCC |  |  |  |  |  |
| --- | --- | --- | --- | --- | --- | --- |
| | Control | 0.01μM | 0.1μM | 1μM | 10μM | 100μM |
| SCPM | $0.97 \pm 0.01$ | $0.96 \pm 0.04$ | $0.96 \pm 0.02$ | $0.97 \pm 0.02$ | $0.89 \pm 0.08$ | $0.88 \pm 0.05$ |
| TCPM | $0.97 \pm 0.01$ | $0.96 \pm 0.03$ | $0.96 \pm 0.02$ | $0.97 \pm 0.02$ | $0.89 \pm 0.07$ | $0.88 \pm 0.05$ |
| FM | $0.97 \pm 0.01$ | $0.96 \pm 0.04$ | $0.96 \pm 0.02$ | $0.97 \pm 0.02$ | $0.89 \pm 0.08$ | $0.89 \pm 0.05$ |
| Inv-SCPM | - | - | - | - | $0.89 \pm 0.08$ | $0.87 \pm 0.05$ |
| Sym-SCPM | - | - | - | - | $0.90 \pm 0.07$ | $0.91 \pm 0.04$ |
| Inv-FM | - | - | - | - | $0.89 \pm 0.08$ | $0.88 \pm 0.05$ |
| Sym-FM | - | - | - | - | $0.14 \pm 0.03$ | $0.06 \pm 0.01$ |

**Figure 4:**
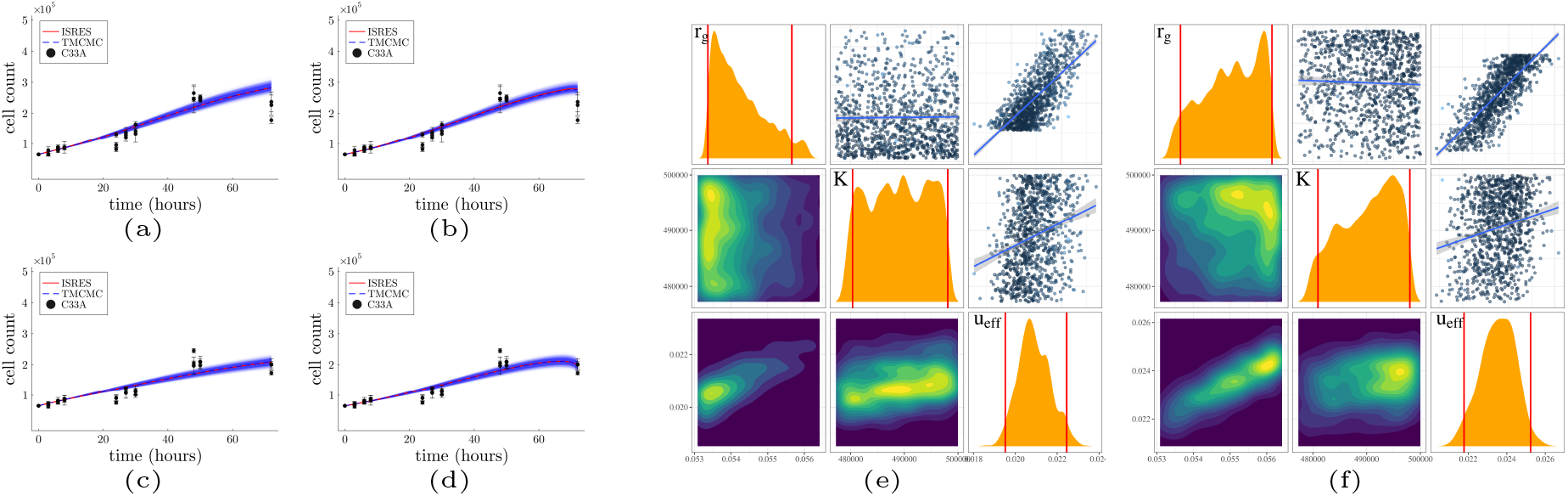
Calibration and uncertainty quantification of the multiscale single cell population models. (a), (b) 10μM, (c), (d) 100μM of 5FU. The model is integrated with (Left column) inverted asymptotic gene expression model, and (Right column) the *y*-symmetric gene expression model. Parameter posterior distributions of the macroscopic model integrated with (g) the inverted gene expression model, (h) the *y*-symmetric gene expression model for the 100μM of 5FU datasets.

**Figure 5:**
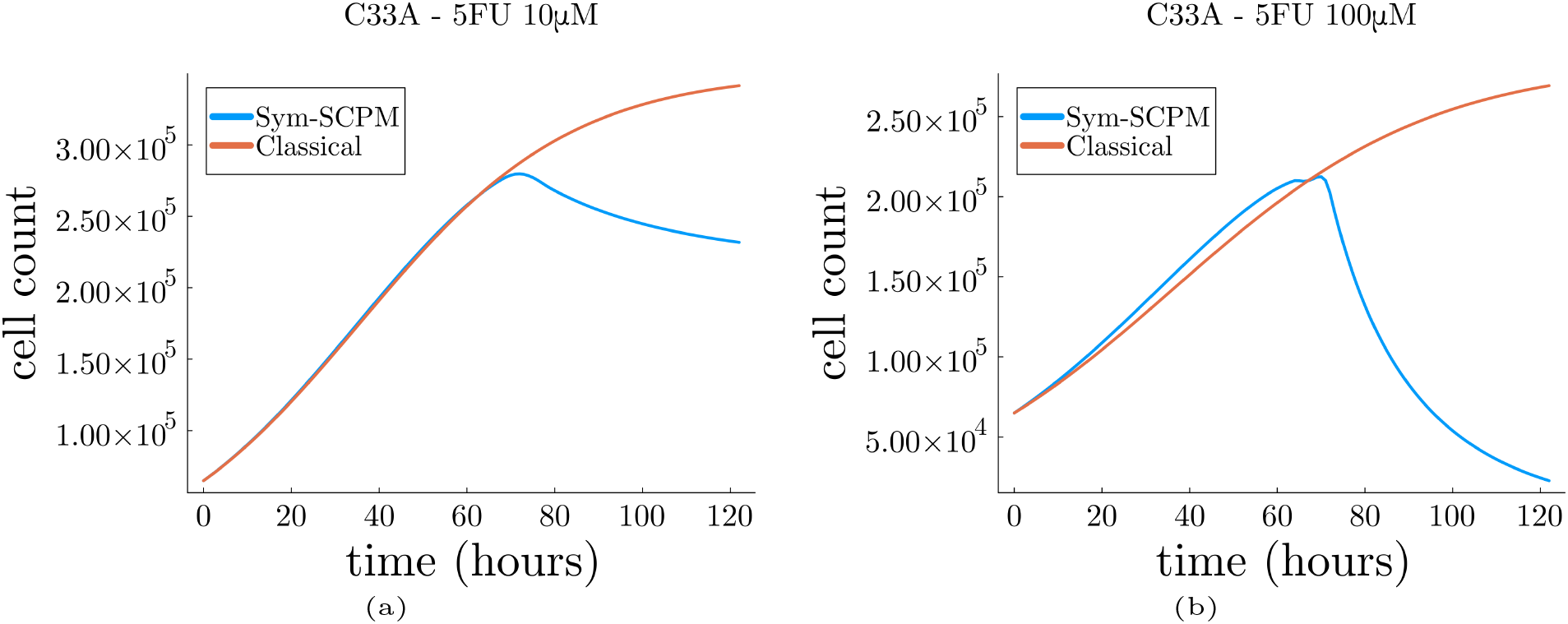
Comparison of dynamics of the classical and the sym-SCPM models with estimated parameters for (a) 10 μM and (b) 100 μM dose of 5FU. The sym-SCPM trajectory is more plausible as we observed cell elimination in the experiments after prolonged times. On the other hand, the classical SCPM predicted stabilisation of the cell population.

### 3.5. Identifiability of the Sym-SCP model parameters

The multiscale Sym-SCP exhibited the best agreement with the experimentally measured cell counts. Here we investigate the identifiability of this model.

#### 3.5.1 Structural identifiability

We demonstrate the structural identifiability of the Sym-SCP model. We start by non-dimensionalizing the model; by setting *n* = *N/K*, the model becomes

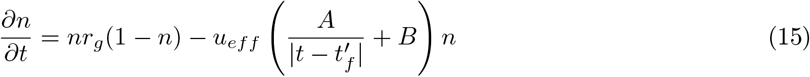

where *t*^′^_*f*_ = *t*_*f*_ + 0.1. For simplicity, we set *r*_*g*_ = *r*, and *u*_*eff*_ = *u*. Since *t*_*f*_ is the final time of the experiment, and 0 ≤ *t* ≤ *t*^′^_*f*_, we can write

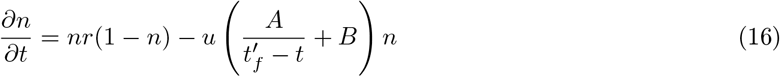

We assumed that all parameters are non-negative. At *t* = 0, we have *n*(0) = *n*_0_, and 0 *< n*_0_ *<* 1 due to the non-dimensionalization. The model can be written in the input-output form

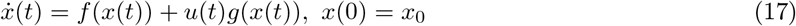

where *f* and *g* are

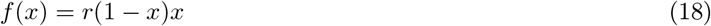

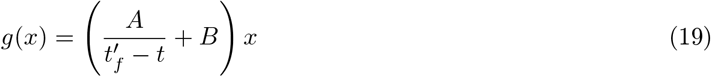

and *x*(*t*) = *n*(*t*). We can write

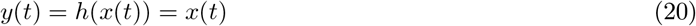

and introduce *p* to be the vector of the parameters to be identified, i.e.

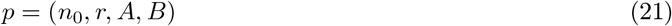

A system as in (17) is uniquely structurally identifiable if the map

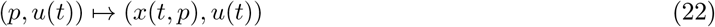

is injective almost everywhere [56, 57].

Assuming *perfect* (continuous and noise-free) input-output data given in the form of *y* and its derivatives at any time interval, we can write the measurements in the form of

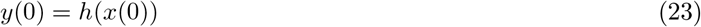

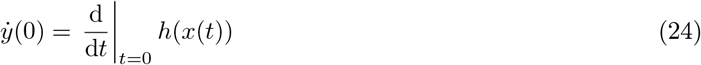

and relate their values to the unknown parameters found in (21). The system is identifiable if there exist inputs, *u*(*t*), such that the system of (23), (24) may be solved for *p*. The right hand sides of (23), (24) can be computed in terms of the Lie derivatives of *f* (18) and *g* (19), found in (17). Recall that Lie differentiation of a C^*ω*^ function H by a C^*ω*^ vector field *X, L*_*X*_ *H*, is

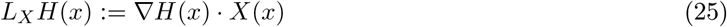

The iterated Lie derivatives are well defined, a nd c an b e w ritten a s *L*_*Y*_ *L* _*X*_ *H* = *L*_*Y*_ (*L*_*X*_ *H*), and 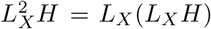.

We can define the observable quantities as the zero-time derivatives of the output *y* = *h*(*x*) as follows

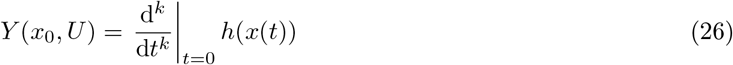

where *U* ∈ ℝ^*k*^ is the control value, *u(t)*, and its derivatives at t = 0; *U* = (*u*(0), *u*^′ (*k*−1)^(0)). The observation space is then defined as the span of the *Y* (*x*_0_, *U*) elements

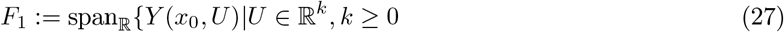

Conversely, the span of iterated Lie derivatives with respect to the output *h* and the vector fields *f, g* is

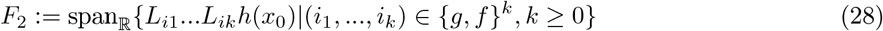

Following [58, 18], *F*_1_ = *F*_2_, hence identifiability may be formulated as the reconstruction of parameter in *p* from elements in *F*_2_. Thus, *p* is identifiable if the map

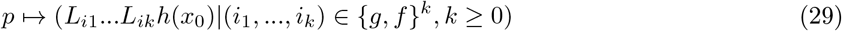

is “1-1”.

Computing the Lie derivatives, we can determine the parameters of *p*

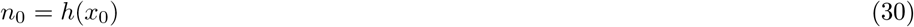

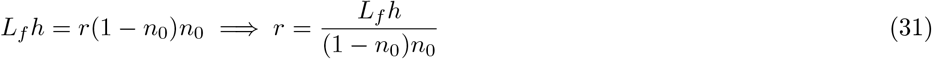

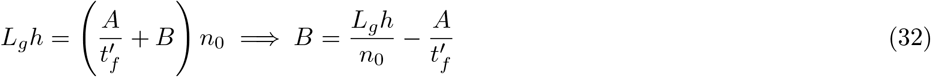

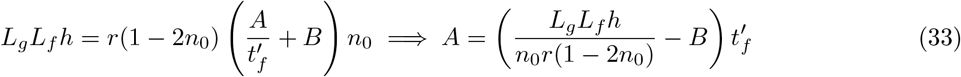

As a result, all parameters of (16) are identifiable.

#### 3.5.2. Practical identifiability

To examine the practical identifiability of the Sym-SCPM we performed the following steps. First, we expanded the prior distribution boundaries of the Sym-SCP model parameters to the values used for the SCP model calibration with the control data. Then, we performed Bayesian inference of the Sym-SCPM using the treatment datasets for 10 and 100 μM FU. The resulting posterior distributions are presented in Fig. 6a and 6b and show unimodal distributions that are significantly narrower than the prior distributions in Fig. 6c. The uncertainty in the output, Fig. 6d and 6e, was increased compared to Fig. 4c and 4d; however, this uncertainty followed the trend of the data and was found within the overall range of the experimental cell counts. Thus, the Sym-SCP model can be considered as practically identifiable for the given dataset.

**Figure 6:**
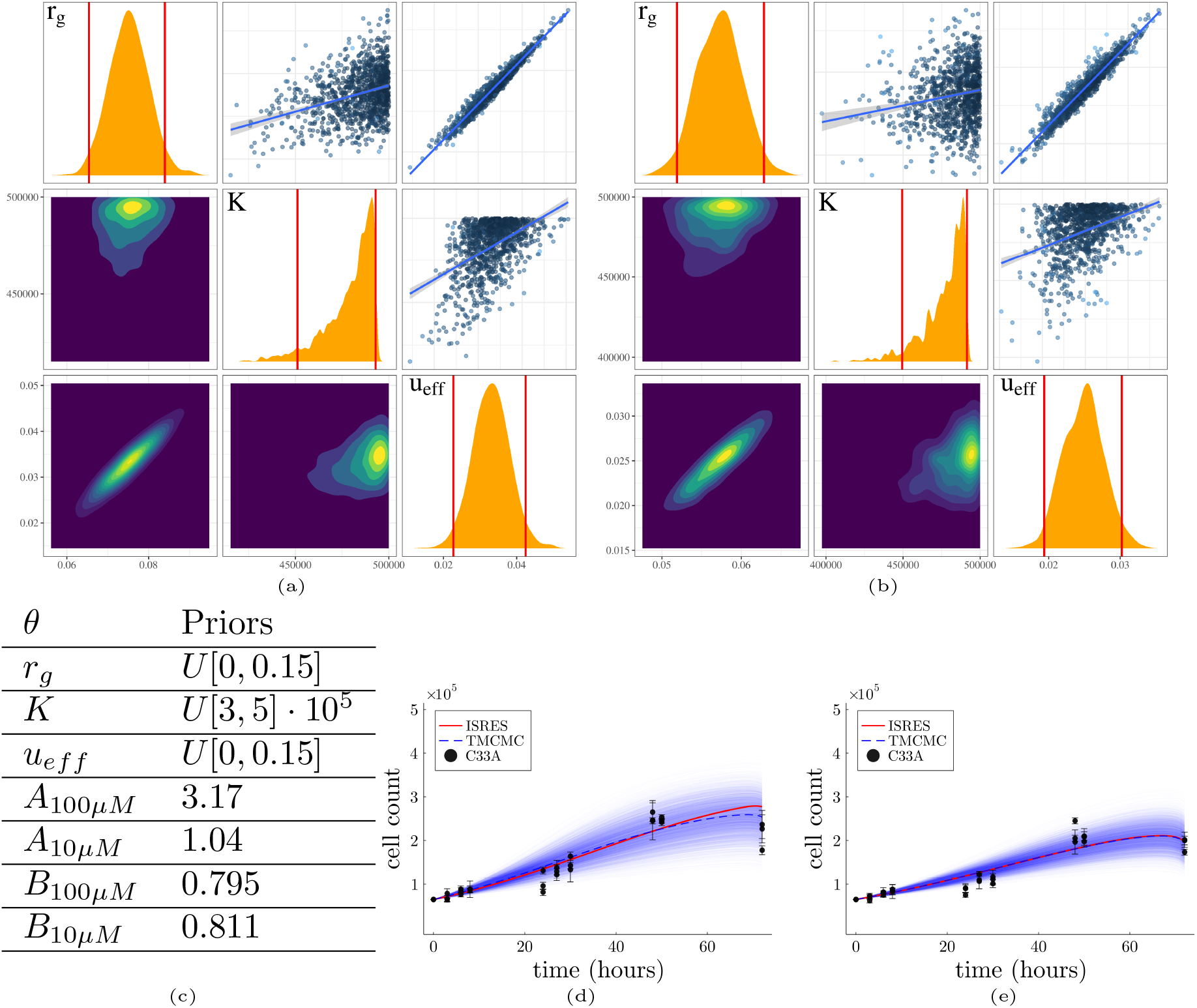
Practical identifiability analysis of the Sym-SCPM with known parameters *A* and *B*. (a), (b) Posterior manifold for the calibration with the 10μM, and 100μM datasets, respectively. (c) Table of the model parameters and their corresponding prior distributions. The *A* and *B* are calibrated from the gene expression data and they are fixed. (d), (e) *In-vitro* and *in-silico* cell counts and their corresponding uncertainties.

#### 3.5.3. The knowledge of gene expression estimates is necessary for a practically identifiable model

The Sym-SCP model was practically identifiable. As shown in Fig. 6c, the parameters *A* and *B* were estimated from the gene expression data and fixed for the identifiability analysis. In this section, we considered the scenario of absence of the gene expression data. Therefore, we considered the values of *A* and *B* to be unknown, and repeated the same process as in the previous section with the prior parameter distributions of Fig. 7c. The resulting posterior distributions, found in Fig. 7a, 7b show increased uncertainty, a result that is reflected to the model output, Fig. 7d, 7e, making the model practically unidentifiable. Thus, the information obtained from the gene expression data for the calibration of the parameters *A* and *B* is necessary for a practically identifiable model.

**Figure 7:**
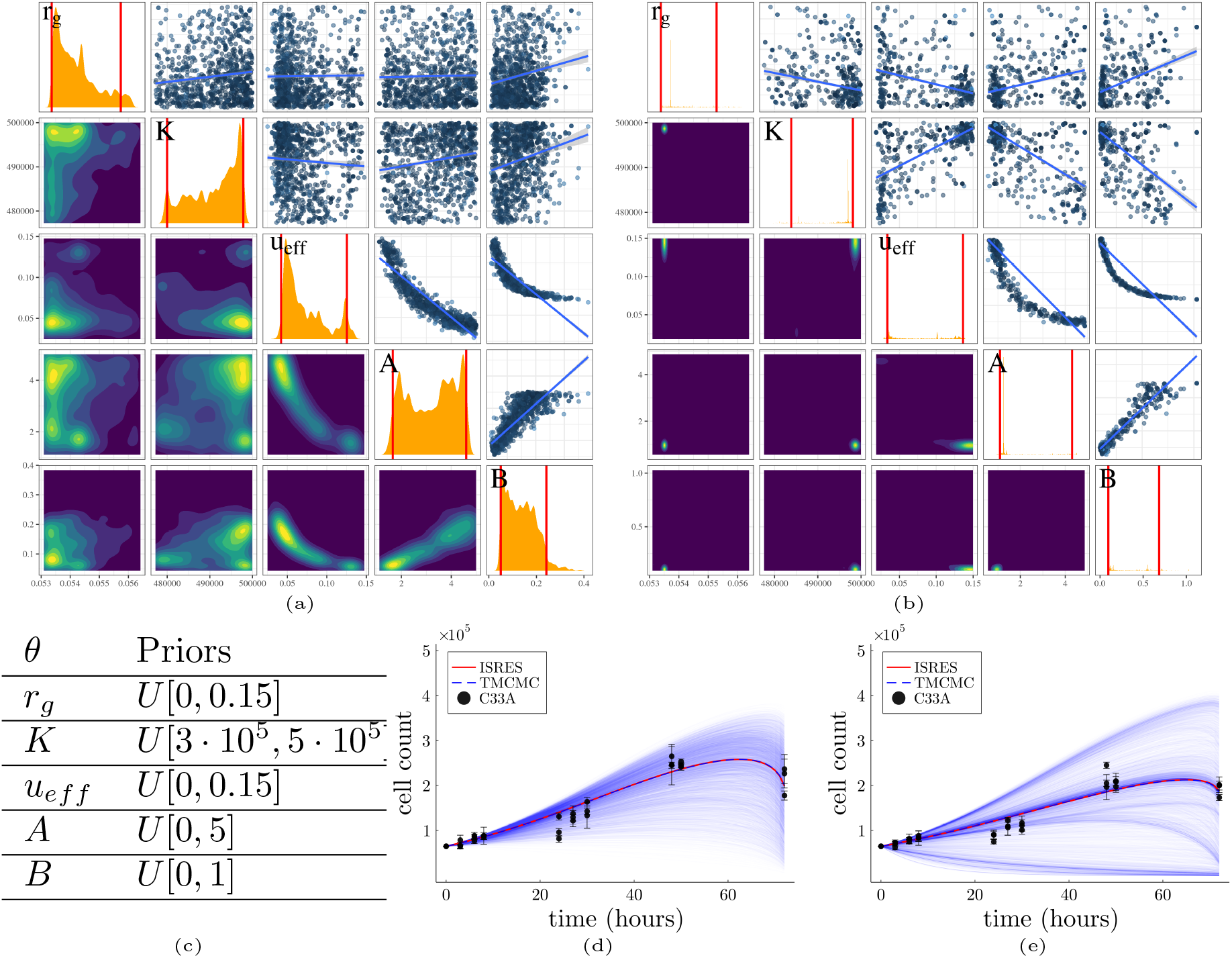
Practical identifiability analysis of the Sym-SCPM considering parameters *A* and *B* unknown. (a), (b) Posterior manifold for the calibration with the 10μM, and 100μM datasets, respectively. (c) Table of the model parameters and their corresponding prior distributions. (d), (e) *In-vitro* and *in silico* cell counts and their corresponding uncertainties. The unknown parameters *A* and *B* increase dramatically the uncertainty of the model output. Thus, the integration of the macroscopic cell count measurements with the gene expression data is necessary for a practically identifiable model.

**Figure A.8:**
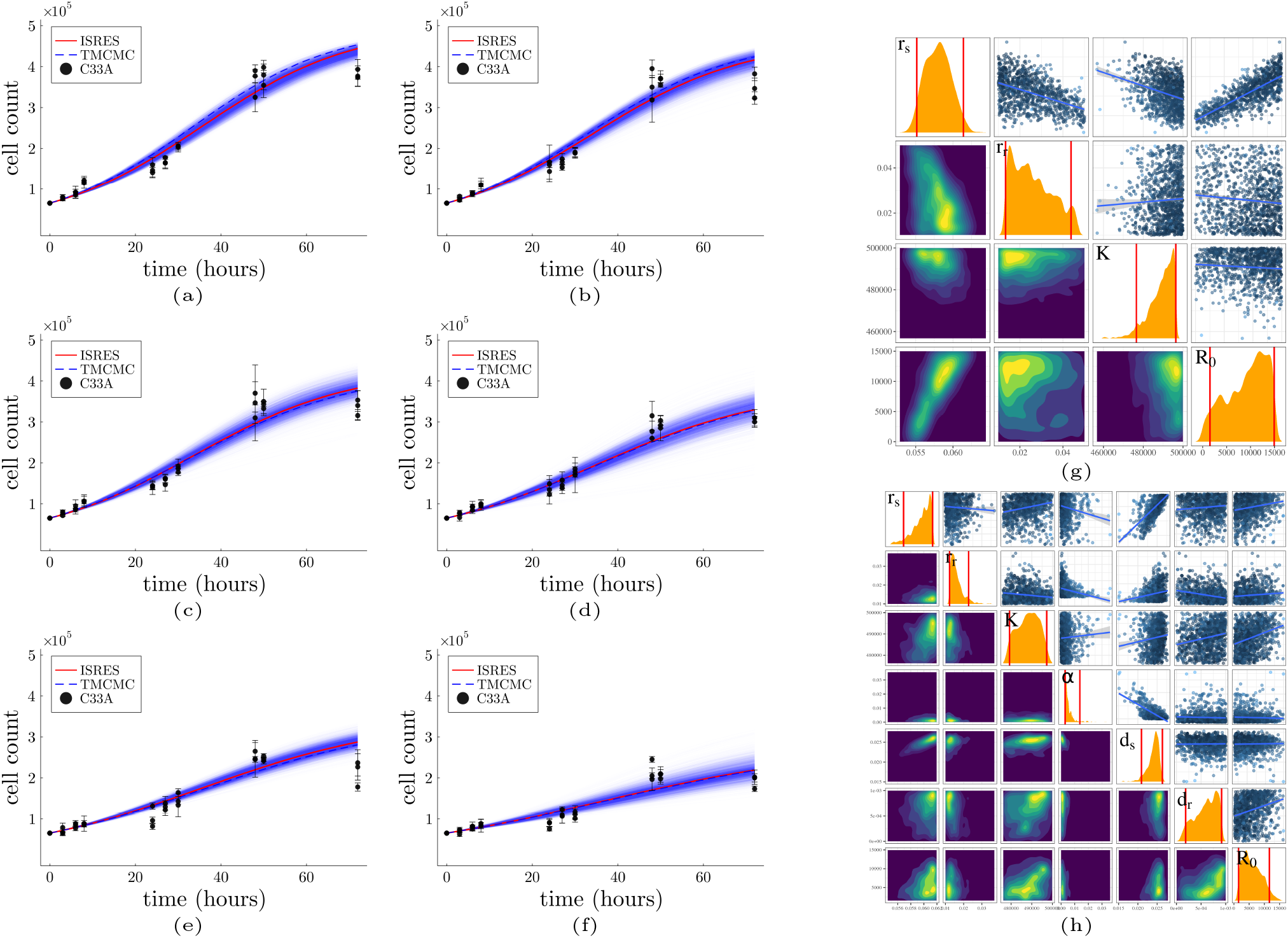
Calibration and uncertainty quantification of the classical two cell population model. (a) Control, (b) 0.01 μM, (c) 0.1 μM, (d) 1 μM, (e) 10 μM, and (f) 100 μM of 5FU datasets. Posterior distributions of the model parameters for (g) control, (h) 100 μM of 5FU datasets. The prior boundaries of the *r*_*s*_, *r*_*r*_ and *K* parameters of the treatment model are extracted from the 95% CI of the posterior distributions of the corresponding parameters of the control model. The red lines denote the 95% CI of the posterior distributions.

**Figure A.9:**
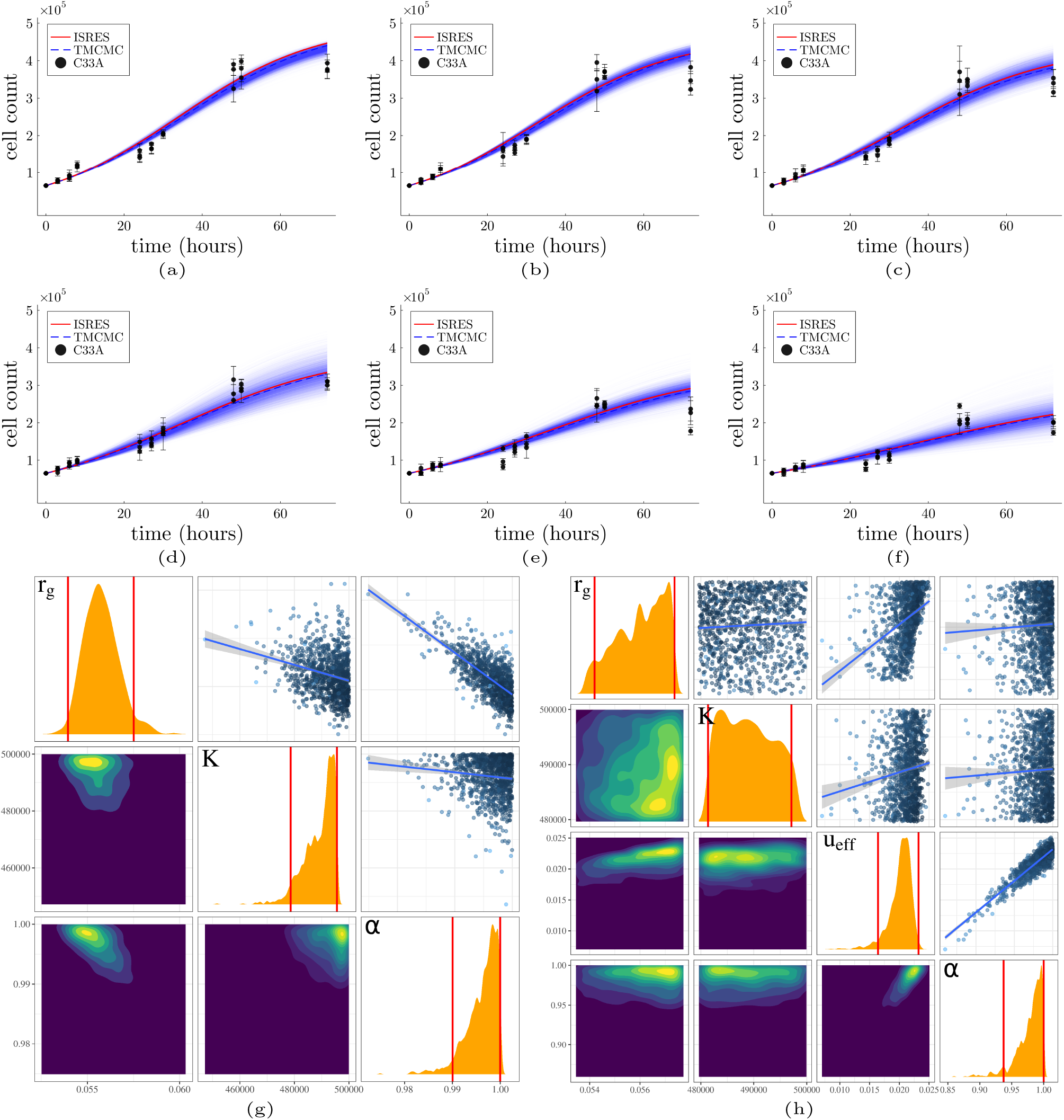
Calibration and uncertainty quantification of the fractional single cell population model. (a) Control, (b) 0.01 μM, (c) 0.1 μM, (d) 1 μM, (e) 10 μM, and (f) 100 μM of 5FU datasets. Posterior distributions of the model parameters for (g) control, (h) 100μM of 5FU datasets. The prior boundaries of the *r*_*g*_ and *K* parameters of the treatment model are extracted from the 95% CI of the posterior distributions of the corresponding parameters of the control model.

**Figure A.10:**
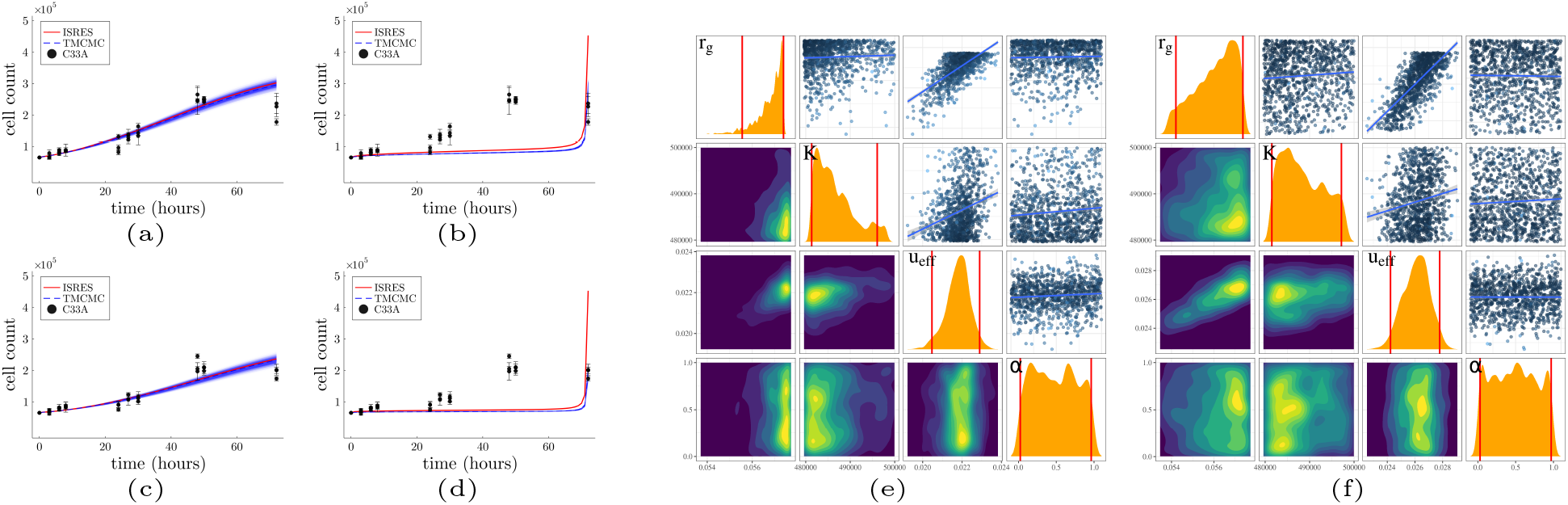
Calibration and uncertainty quantification of the multiscale fractional single cell population models. (a), (b) 10 μM, (c), (d) 100 μM of 5FU datasets. The model is integrated with (Left column) inverted asymptotic gene expression model, and (Right column) the *y*-symmetric gene expression model. Parameter posterior distributions of the macroscopic model integrated with (g) the inverted gene expression model, (h) the *y*-symmetric gene expression model for the 100μM of 5FU datasets.

## 4. Discussion

In this study, we examined the predictive performance of a variety of mathematical models describing *in vitro* cervical cancer growth and its response to 5FU treatment. In addition, we investigated the integration of cell counts and OCT4 expression into multiscale models to improve predictive performance. The analysis yielded important insights into the agreement of the calibrated models with the experimental data and their uncertainty, both in terms of parameters and final output. In this section, we discuss the fitting performance of the examined models, the performance of the two calibration methods, and the identifiability of these models.

### 4.1. Fitness and Interpretation of model behaviour

We investigated the behaviour of 7 mathematical models of tumour growth in relation to longitudinal *in-vitro* measurements of tumour growth and the expression of genes associated with changes in chemosensitivity. The macroscopic SCPM exhibited good agreement with the data, especially for the no-treatment and low-dose conditions, and a low uncertainty in its output prediction (Fig. 2). The agreement with the experiments decreased for the two highest doses of 5FU, with the model exhibiting a behaviour that was more linear rather than sigmoidal. This behaviour can be explained by the carrying capacity compared to the maximum number of cells found in the cell cultures. The carrying capacity was estimated by the control data and was significantly higher than the cell counts found in the high dose treatment conditions. Therefore, it was not possible to reach a plateau at later time points.

To improve accuracy at the highest doses, we expanded our analysis to examine more complex models. Specifically, we considered a TCPM that takes into account drug-sensitive and drug-resistant cell populations. The resulting agreement with the experiments did not improve, and the uncertainty of the output increased slightly (Fig. A.8). In addition, the initial number of drug-resistant cells remained highly uncertain for both the control and treatment data, probably due to their low growth rate. The results showed that TCPM did not improve the agreement or uncertainty and the obtained results do not unequivocally suggest the emergence of drug resistance.

Fractional calculus is a relatively new area of mathematics that allows the ordinary integer order of derivatives to assume non-integer values. Fractional derivative models can take into account non-local interactions in space and dynamic memory in the time domain [59]. In this context, we considered a SCPM with a non-integer order of the time derivative and treated the derivative order as unknown, leading to the corresponding FM. The calibration of the FM yielded a derivative order close to 1, suggesting that static fractional derivatives did not improve the performance of the SCPM (Fig. A.9).

### 4.2. Evidence for the use of multiscale models and hypotheses

Multiscale expansion of the SCPM and FM was achieved by constructing an expression model for OCT4 and relating it to cell death rates (Fig. 3). The two cell death rate functions were coupled with the cell death term of the SCPM and the derivative order of the FM, resulting in Inv-SCPM, Sym-SCPM, Inv-FM, and Sym-FM respectively. Sym-SCPM showed the best agreement with the experimental data for the highest doses (10 μM and 100 μM 5FU) and low uncertainty in the output, resulting in a more accurate dynamic trajectory than the other models (Fig. 4, 5). The increase in the agreement can be explained by the fact that the cell death rate exhibited a steep increase during the later days of the experiment. In turn, this suggests that there is a threshold level of OCT4 expression that triggers or suppresses cell death (i.e. gainof-function and loss-of-function, respectively) and that OCT4 may activate or suppress mechanisms related to chemoresistance as a binary function rather than a continuous function of time.

The form of the Sym-SCPM is not arbitrary; it was based on observations of growth curves, gene expression estimates, and recent findings from the literature [34, 35, 36]. From the growth curves, we observed a sigmoid trend in the presence and absence of therapy, even though the treatment was initiated at *t* = 0 h. Instead, we would expect the cell counts to stabilize or decrease in the presence of the treatment. This observation led us to hypothesize that the cell death rate was not static and may exhibit a delayed increase after treatment initiation. At the same time, we observed that the expression of OCT4 – a potential regulator of chemosensitivity in cells [34, 35, 36] – decreased over time. In addition, studies in bacteria [29, 30] have shown that there is a dependence between gene expression rates and growth rates. We therefore hypothesized that the cell death rate is linked to the expression of OCT4 and investigated the two possible transient cell death rates.

On the other hand, Sym-FM exhibited the worst performance compared to the other models. This is justified by the fact that the derivative order remains low most of the time, resulting in a time delay of the model’s entry into the exponential phase. The sudden increase in the derivative order at later time-points leads to a rapid entry into the exponential phase, which is inconsistent with the experimental observations. In addition, the scaling parameter of the derivative order *α* remains highly insensitive to the model output. This result suggests that the overall dynamic behaviour of the derivative order seems to be more important than its amplitude.

### 4.3. Structural and practical identifiability of the Sym-SCPM

The Sym-SCPM yielded the best performance during high dose treatment conditions. Nevertheless, since this model contains a novel cell death term, it was necessary to further investigate the identifiability of the model parameters. This allowed us to examine whether the model parameters can be determined by the model outputs (structural identifiability) and estimate their uncertainties given the available data (practical identifiability) [39]. The structural identifiability analysis was performed using the methodology presented in [52]. The Sym-SCP model, which included both the macroscopic parameters and the parameters corresponding to transient cell death, was found to be structurally identifiable. The practical identifiability analysis was performed using Bayesian inference, using the same prior distributions as in the SCP model calibrated with the control data. The model was found to be practically identifiable, provided that the parameters of the transient cell death term were known. These two parameters were estimated from OCT4 expression data (Fig. 6). The model did not pass the identifiability test for completely unknown parameters (Fig. 7), suggesting that information on gene expression levels is crucial.

### 4.4. Monte Carlo and Global optimization algorithms

We used two calibration methods in this study; the likelihood optimization method using the ISRES algorithm, and Bayesian inference using the TMCMC algorithm. The ISRES algorithm is considered to be a global or semiglobal optimization method. It uses a population size of 20 × (*n*+1) candidate solutions, where *n* is the number of parameters. In contrast, TMCMC used 1024 candidate solutions for each generation. Overall, the two methods showed comparable performance in terms of finding the maximum likelihood, and TMCMC was able to find parameters reuslted in a model that achieved better performance than ISRES. Although the ISRES algorithm, and likelihood function optimization methods in general, are considerably faster than Bayesian inference and Monte Carlo methods, recent efforts to parallelize of the MCMC algorithm have significantly improved its efficiency. The TMCMC algorithm is inherently parallel, and its function does not require a burn-in period such as traditional MCMC methods. In our case, the TMCMC algorithm took about 10 minutes to run on a 4 × Intel Core™ i7-6820HQ CPUs running at 2.70GHz. Overall, Bayesian inference and the TMCMC algorithm provide information not only on the fitness of the model, but on its uncertainty and identifiability. Together with its improved performance, it can be considered as an efficient all-in-one tool.

## 5. Conclusions

We performed a rigorous calibration and uncertainty quantification analysis of classical, fractional, and multiscale models of tumour growth using data obtained from cell cultures of the C33A cell line during control and therapy conditions. We also provided a method of integrating information coming from different levels into a multiscale modelling framework, which allowed us to obtain more accurate model predictions in the presence of sparse time-course tumour growth measurements. Specifically, our results suggest that the single-cell population model was sufficient for describing tumour growth under no-treatment and low dose 5FU treatment conditions. They also showed that the multiscale expansion of this model by using data on the expression of the transcription factor OCT4 improved the agreement with the experimental observations for the two highest doses of 5FU. Investigation of two potential mechanisms for the transient cell death rate indicated that the cell death process is more likely to function as a binary mechanism triggered above a threshold level of OCT4 expression rather than a more gradual activation inversely related to OCT4 expression. The two-cell population model and fractional models did not significantly improve performance. The Sym-SCPM proved to be both structurally and practically identifiable, suggesting that its parameters can be unambiguously estimated. Overall, we anticipate that the proposed methodology will enable researchers to design more accurate models, as sparse multilevel data are available in most cancer research studies. The data and code associated with this work can be found at https://github.com/NMDimitriou/ModTherapy

## Appendix

A. Analytical solution of the fractional model

To solve the fractional model, first we need to provide some useful definitions obtained from [59, 60].

### Definition 1

Let *n* ∈ ℕand *ν* a non-integer number with Re(*ν*) *>* 0, the Gel’fand-Shilov function is defined as

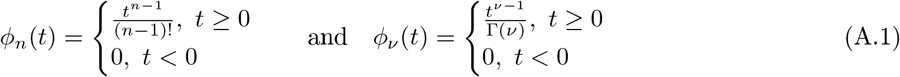

### Definition 2

Let *α >* 0, the Mittag-Leffler function is defined by

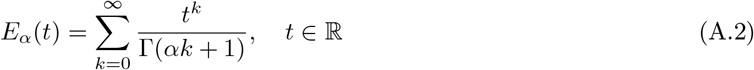

The two-parameter Mittag-Leffler function with *α, β >* 0 is defined by

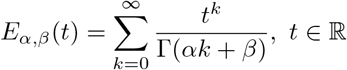

### Definition 3

Let *n* ∈ ℕand *f* (*t*) : ℝ → ℝ an integrable function. The Integral Operator *I* of order 1 and *n* is defined by

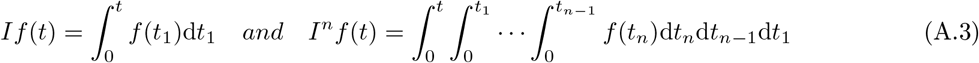

### Theorem 1

*For f* (*t*) : ℝ →, *the n-order integral of f will be*

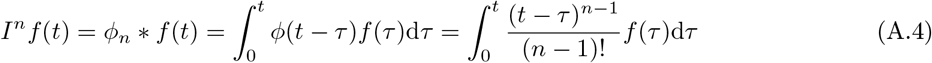

*where* ##x002A; *denotes the Laplace convolution product, and ϕ*_*n*_(*t*) *is the Gel’fand-Shilov function*.

∈

### Definition 4

Let *f* (*t*) be an integrable function and *ν* ∈ ℂ, such that Re(*ν*) *>* 0, the Riemann-Liouville fractional integral of order *ν* of *f* (*t*), denoted by *I*^*ν*^ *f* (*t*), is defined as

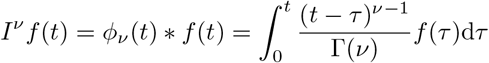

### Definition 5

Let be a differentiable function, *m* ∈ ℕ and α ∉ ℕ such that *m* − 1 < Re(*α*) < m. Caputo’s fractional derivative is defined by

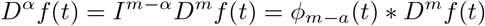

For the solution of the fractional logistic model we need the laplace transform of Caputo’s derivative that is

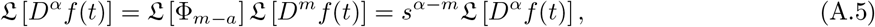

and of the following expression involving the Mittag-Leffler function

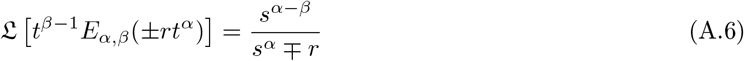

Additionally, an important relation between the two-parameter and one-parameter Mittag-Leffler functions is given by

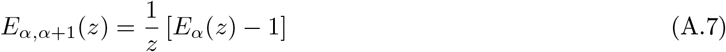

Taking the classical logistic model, and setting *u* = 1*/N* we get

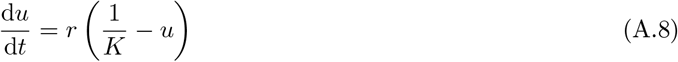

Thus, the fractional version of the model can be written as

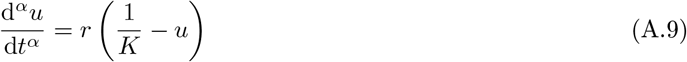

where the derivative order is 0 *< α* ≤ 1. Applying the Laplace transform to (A.9) we get

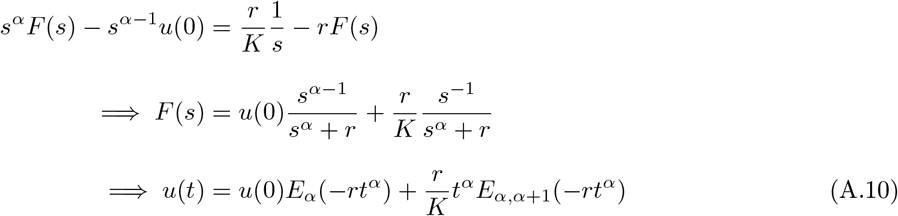

Using the relation found in (A.7), we get

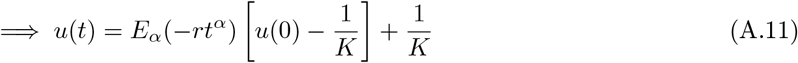

Setting *N* (*t*) = 1*/u* results

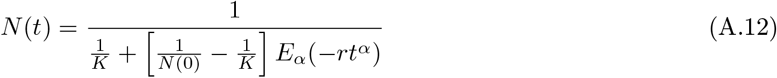

Similarly, by incorporating the therapeutic term in (A.9) we get,

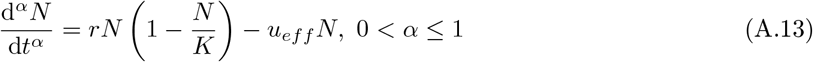

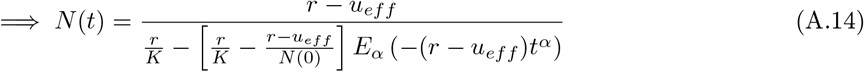

## Notes

https://github.com/NMDimitriou/ModTherapy

### Competing Interest Statement

The authors have declared no competing interest.

### Summary of Updates

Corrected typos, moved 2 figures in the appendix.

